# Towards machine learning fairness in classifying multicategory causes of deaths in colorectal or lung cancer patients

**DOI:** 10.1101/2025.02.14.638368

**Authors:** Catherine H. Feng, Fei Deng, Mary L. Disis, Nan Gao, Lanjing Zhang

## Abstract

Classification of patient multicategory survival outcomes is important for personalized cancer treatments. Machine Learning (ML) algorithms have increasingly been used to inform healthcare decisions, but these models are vulnerable to biases in data collection and algorithm creation. ML models have previously been shown to exhibit racial bias, but their fairness towards patients from different age and sex groups have yet to be studied. Therefore, we compared the multimetric performances of 5 ML models (random forests, multinomial logistic regression, linear support vector classifier, linear discriminant analysis, and multilayer perceptron) when classifying colorectal cancer patients (*n*=515) of various age, sex, and racial groups using the TCGA data. All five models exhibited biases for these sociodemographic groups. We then repeated the same process on lung adenocarcinoma (*n*=589) to validate our findings. Surprisingly, most models tended to perform more poorly overall for the largest sociodemographic groups. Methods to optimize model performance, including testing the model on merged age, sex, or racial groups, and creating a model trained on and used for an individual or merged sociodemographic group, show potential to reduce disparities in model performance for different groups. Notably, these methods may be used to improve ML fairness while avoiding penalizing the model for exhibiting bias and thus sacrificing overall performance.

## Introduction

Artificial Intelligence (AI) is changing medicine, with growing applications of Machine Learning (ML) algorithms in the biomedical field, and more specifically in cancer research and treatment [1-3]. Lung and colorectal cancers (CRC) are the two most common causes of cancer deaths in the United States[4]. As various types of ‘omics data, including transcriptomic and genomic data, have been generated in recent decades[3], ML algorithms have been used to predict disease prognosis and helped inform patient treatments for lung cancers[5-8] and CRC[9-11]. However, these models are vulnerable to biases in both data collection and algorithm creation, often resulting in minority groups such race- and age-minorities being fit into a model designed for the majority[12-17].

Racial biases have been shown in ML models used in US healthcare systems, where the model underperforms for certain race group(s)[18-20]. However, accuracy alone was considered when evaluating the fairness of the model. We have previously shown that although accuracy is important, additional metrics such as precision, recall, and F1 may help to better understand the model’s performance and thus evaluate its fairness[5, 10, 21]. In addition, while racial biases were considered, studying ML model performance with different age and gender groups can further inform us about other biases within these models as reported in non-cancer medical field[17, 22-25]. The existence of these biases warrants better understanding of which algorithms exhibit the least bias towards patients from different sociodemographic groups, and whether these algorithms can be optimized to improve model fairness. Whereas previous studies have investigated this question, many penalize ML models for exhibiting bias, reducing bias but at the cost of overall performance[26-28]. Hence, we aimed to understand the multimetric performance of random forests (RF), multinomial logistic regression (MLogit), linear support vector classifier (Linear SVC), linear discriminant analysis (LDA), and multilayer perceptron (MLP) when classifying four-category outcomes for CRC and lung adenocarcinoma patients from various sociodemographic groups. We also aimed to reduce bias of model performances, including testing the model on merged age, sex, and racial groups, and training and testing the model on an individual or merged sociodemographic group. We found that both methods held the potential to increase algorithmic fairness (i.e., reduce biases) for certain ML models, and that the latter method can do so with significantly reducing overall ML performance.

## Methods

### Data Extraction

We obtained individual-level data of colorectal and lung adenocarcinoma data from The Cancer Genome Atlas Program (TCGA) Pancancer Atlas through the cBioPortal repository[29], as described before[5, 7, 9, 10, 21]. The data were publicly available and de-identified, making this an exempt study (category 4) that did not require an Institutional Review Board IRB review.

We performed differentially expressed genes (DEG) analysis to remove the less relevant genes using Chi-square test. The outcomes of the classification models were the patients’ 4-category survival. This included Alive with No Known Disease Progression, Alive with Disease Progression, Dead with No Known Disease Progression, and Dead with Disease Progression.

We grouped the patients based on their age, race, and sex, the three sociodemographic factors of our interest, to analyze the extent of the biases that ML models had on each group. CRC patients were divided into 3 age groups based on whether they were 33 – 60, 60 – 72, or 73+ years old, and into 5 racial groups that included Race Unknown, American Indian or Alaska Native, Black or African American, White, and Asian and Pacific Islander (AAPI). Meanwhile, Lung patients were split into 2 age groups based on whether they were below 65 years old or 65 and above and into 3 racial groups of Black, White, or Other. Both CRC and Lung patients were also divided into Male and Female groups based on their sex.

### Pipelines

All ML processes were conducted using Python 3.6.9 in Google Colaboratory. We used five ML models, namely RF, MLogit, Linear SVC, LDA, and MLP. For each ML model, we created a pipeline that automated the process of creating, tuning, and collecting the performance metrics Accuracy, Precision, Recall, and F1. Performance metrics were obtained using the cross_validate function imported from the sklearn.model_selection package from Scikit Learn. All tuned models were run 20 times to obtain the mean and standard deviation of each performance metric.

As **Figure 1** shows, we applied the ML models that were tuned and trained on all the patients to each group (row A). We also used these models to examine whether merging groups would improve differences between or among the groups (row B). Finally, we applied ML models that were trained on each patient or merged group to that same group (row C). All processes were first performed on the CRC dataset before being repeated in the Lung dataset to validate our findings.

**Figure 1.**
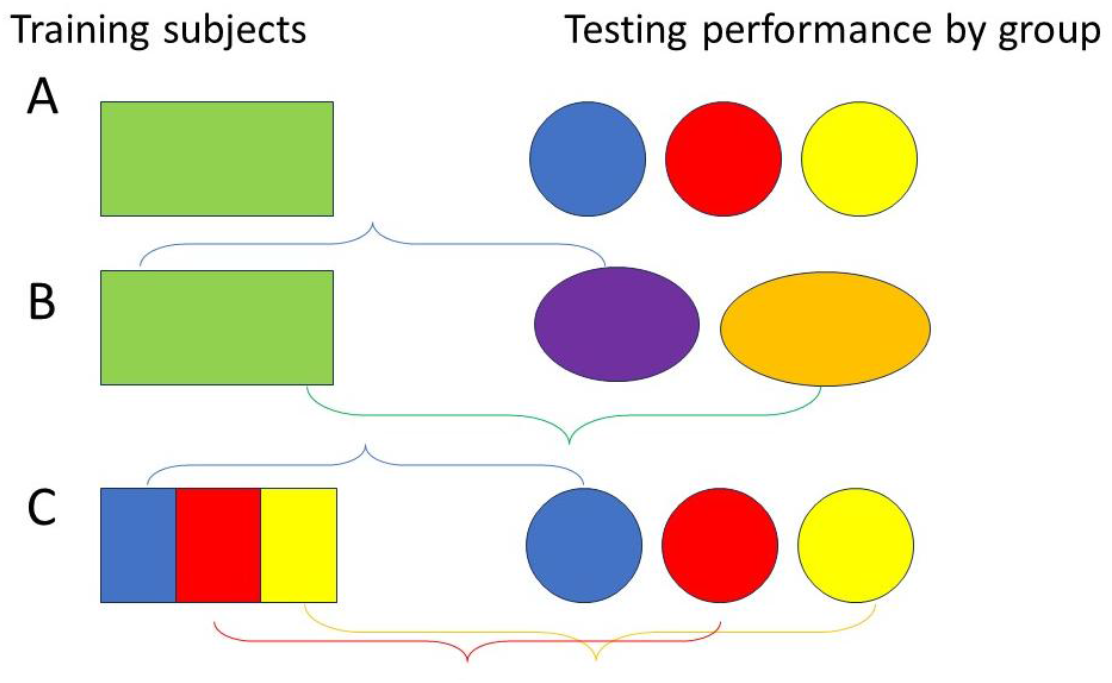
Three grouping approaches to improve machine learning fairness and reduce biases.

### Comparing Different Models

To examine whether, and to what extent, biases in our sociodemographic factors of interest impacted model performance, we first tuned and trained the models on the entire dataset before testing the models on each individual age, race, and sex group.

Because of the small size of the American Indian and AAPI groups in the CRC dataset (1 and 12 patients, respectively), we focused on the Race Unknown, White, and Black or African American patient groups to measure model bias and only included American Indian and AAPI patients when merging groups. Results from each ML model were also compared to examine whether certain models were less affected by biases and to compare the results from different types of models.

For the RF model, we used the RandomForestClassifier from the Python Scikit Learn package and set our split criterion as the Gini impurity index. We automated the tuning process, tuning parameters n_estimator (the number of trees in the forest) and min_samples_split (the minimum number of samples required to split an internal node). Varying n_estimator (ranging from 5 to 145, with intervals of 5) and min_samples_split (ranging from 2 to 11, with intervals of 3), we performed 5-fold cross validation using cross_val_score to collect accuracy values from the Scikit Learn package. The model was tuned three times before the pipeline automatically selected the parameters that produced the highest average accuracy. We then tested the tuned model on individual age, race, and sex groups and collected performance metrics by the group.

For Mlogit modelling, we used LogisticRegression from sklearn.linear_model to perform MLogit and set the solver to newton-cg. We tested the model on individual age, race, and sex groups after training the model on the rest of the dataset to prevent model overfitting. Since we don’t need to tune MLogit, we used KFold from the sklearn.model_selection package to perform 10-fold cross validation before extracting performance metrics.

To create our Linear SVC, LDA, and supervised neural network models, we used SVC from sklearn.svm, LinearDiscriminantAnalysis from the sklearn.discriminant_analysis package, and MLPClassifier from sklearn.neural_network, respectively. The solver aam was used for the MLP model. We split the dataset into training and testing groups using train_test_split from sklearn.model_selection. To test the performance of the tuned model on individual sociodemographic groups, we dropped members of other groups from the aforementioned testing group. For this model, we also used KFold to perform 10-fold cross validations and obtain the model’s performance metrics.

### Combining Groups

We combined individual race and age groups in the CRC and Lung datasets and tested the ML models on these subgroups to test whether that would reduce the gap between model performance for different subgroups.

For CRC, we created subgroups of patients 33-72, 61+, and between 33-60 or 73+ years of age. We also created race subgroups of Unknown/Nonwhite (Race Unknown, Black or African American, American Indian, and AAPI), Race Known (White, Black or African American, American Indian, and AAPI), White/Unknown, and Minorities (Black or African American, American Indian, and AAPI) for the CRC dataset. For Lung, we grouped patients into subgroups of Minorities (Black and Other), Black/White, and Nonblack (White and Other) based on their race.

By creating the subgroups, we effectively reduced the number of groups being compared at a time from 3 or 5 (CRC Age and Lung Race or CRC Race) into 2 subgroups. We tested the models tuned on the entire dataset on these subgroups, using the same automated pipelines as before. We then compared the performance metrics for each subgroup against the remaining individual groups (or subgroups composed of those individual groups) to identify whether differences in model performance across different race and age groups had been reduced or exacerbated.

### Tuning Individual Groups

Finally, we tuned and trained each ML model on individual race, age, and sex groups, as well as the subgroups with combined race and age groups that we created in the last section to identify whether individualizing models for specific groups would decrease ML model biases. The same pipelines and processes as the ones used in the first method (comparing different ML models) were employed here.

Each race, age, and sex group and subgroup was split into training and testing groups. The model was first tuned and trained on the individual group or subgroup before getting tested and performance metrics obtained. These groups were shuffled in the 20 iterations that we performed.

### Statistics and reproducibility

Mean squared error (MSE) was calculated to compare ML performance differences among groups, while standard deviation (SD) was shown the figures. The number of independent experiments, the number of events, and information about the statistical details and methods are indicated in the relevant figure legends. The Student *t* test was used to compare ML performance metrics. P values of less than 0.05 were considered significant.

## Results

### Baseline Characteristics

Using the Chi-square test, we identified 2,034 DEG in the CRC dataset and 2,647 DEG in the Luad one. There were 589 qualified CRC cases, with 406 (68.7%) patients alive with no progression, 65 (11.0%) alive with progression, 35 (5.9%) dead with no known progression, and 85 (14.4%) dead with disease progression **(Supplementary Table 1)** [10]. There were 188 (31.9%) cases between 33 and 60 years of age, 197 (33.4%) between 61 and 72, and 206 (35.0%) that were 73 or older. 310 (52.6%) of the patients were male and 279 (47.4%) female. In addition, the race of 229 (38.9%) cases were unknown. 1 (0.02%) patient was American Native or Alaskan Indian, 64 (10.9%) were Black, 283 (48.0%) were White, and 12 (2.0%) were AAPI.

In the Lung dataset, there were 515 qualified cases, with 252 (48.9%) alive without disease, 76 (14.8%) alive with disease, 80 (15.5%) dead with no known disease, and 107 (20.8%) dead with disease **(Supplementary Table 2)**[7]. 220 (42.7%) cases involved patients under 65 years old and 295 (57.3%) cases were 65 and above. 277 (53.8%) of the patients were male and 238 (46.2%) female. In addition, 52 (10.1%) were Black, 388 (75.3%) were White, and 75 (14.6%) were of another race.

The breakdown of each race, age, and sex group into the four survival categories is shown in Supplementary Tables 1 and 2 as well.

### Model Performance

For all five ML models, the 20 iterations we ran appeared to perform similarly. We found that in most of the models, larger sociodemographic groups tended to perform more poorly overall than smaller ones.

Out of the three CRC age groups, RF, Linear SVC, LDA, and MLP had the highest accuracy for the second largest group, with patients aged 61-72 (73%, 73%, 83%, and 72%, respectively), with LDA having the best overall performance with a precision of 54%, a recall value of 62%, and an F1 value of 68% **(Figure 2)**. Interestingly, the below 65 Lung age group also had the best accuracy among every model except Linear SVC and the best overall performance using MLP (with a 57% accuracy 57%, 41% precision, 45% recall, and 40% F1), despite this group being smaller than the other one **(Figure 3)**.

**Figure 2.**
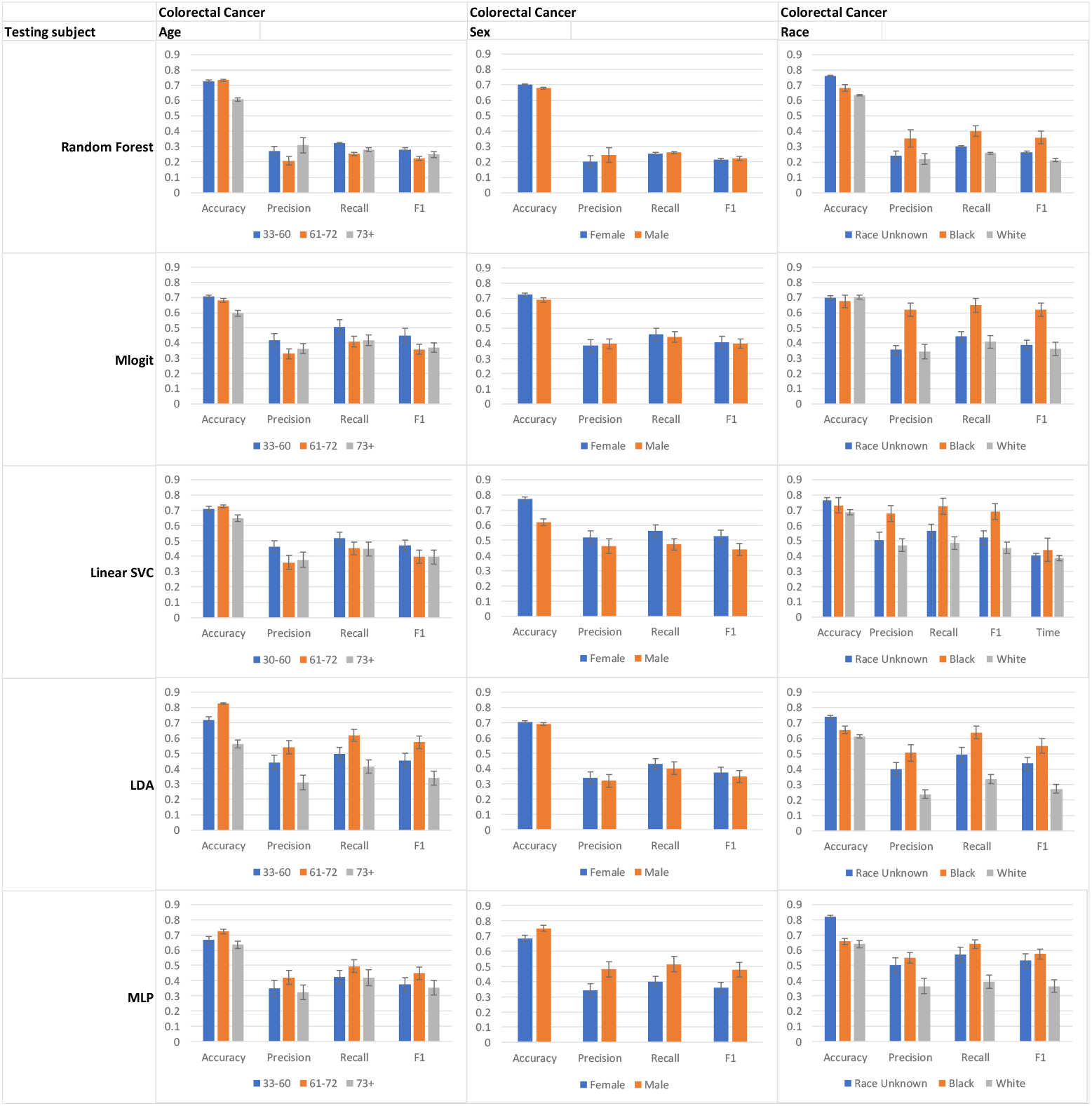
The five ML models’ performance classifying patients in different age, race, and sex groups into their 4-category survival outcomes in the CRC dataset after being trained on patients from each group. Error bar, standard deviation.

**Figure 3.**
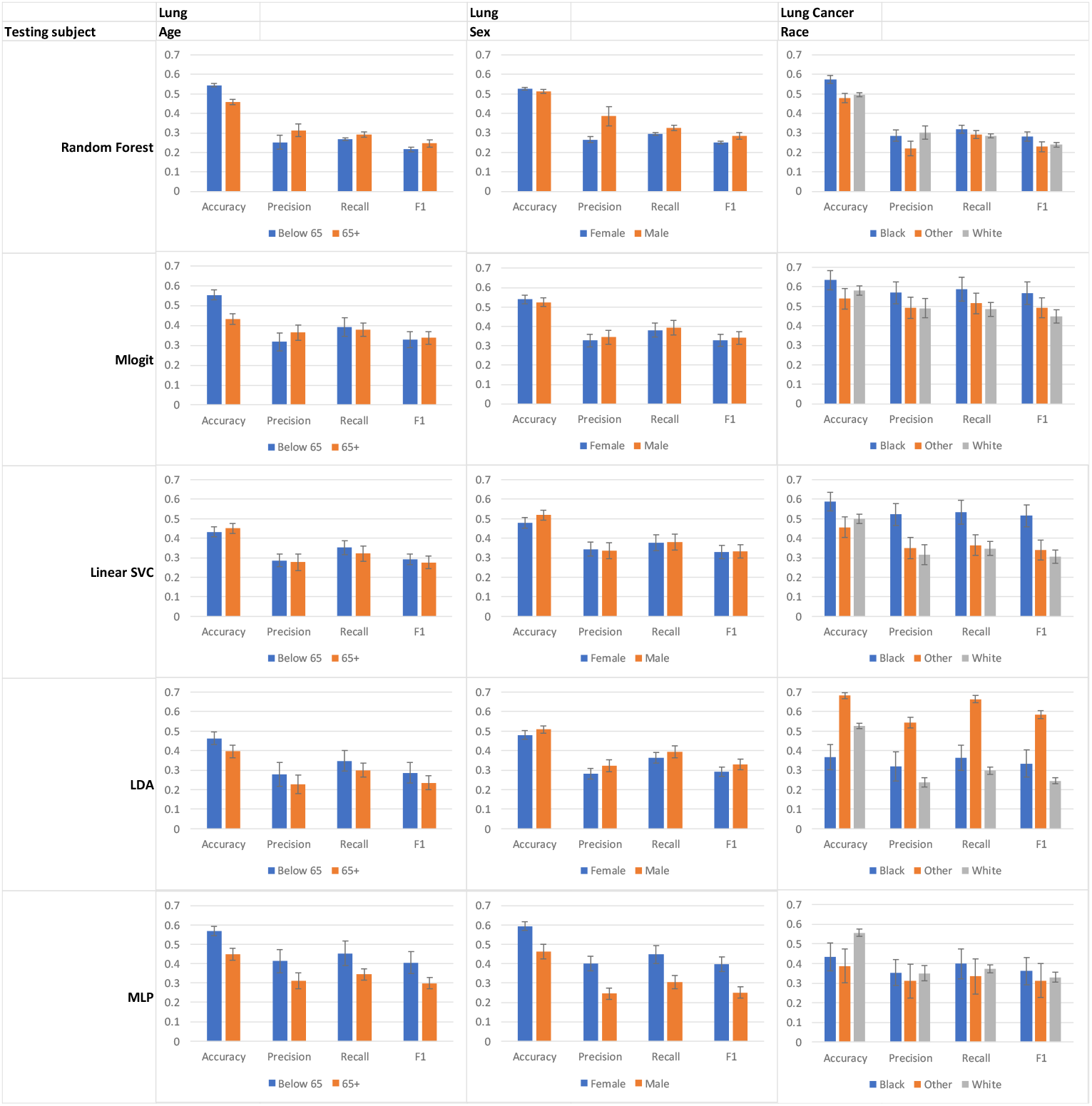
Model performance for lung cancer patients in different sociodemographic groups after being trained on the entire dataset. Error bar, standard deviation.

Both the CRC and Lung datasets had fewer female cases than male. However, the models generally had a higher accuracy for female patients (except for MLP in CRC and Linear SVC and LDA in Lung). The most striking difference in accuracy occurred in Linear SVC for CRC (77% vs. 62%) and MLP for Lung (59% vs. 46%), both of which had the best overall performance for female patients (**Figure 2 & 3)**.

Finally, in CRC race, the second largest group, patients whose race was unknown, typically had the highest accuracy. Strikingly, Black or African American, the smallest group we focused on when comparing different models, consistently had the highest Precision, Recall, and F1 values in all five models, despite not having the highest accuracy. Similar findings were observed in the Lung dataset, where the two smallest groups, Black patients and patients of another race, had the highest Precision, Recall, and F1 values across all of the ML models. However, interestingly, they also had the highest accuracy values in every model except for MLP, unlike what occurred with the CRC racial groups.

While combining individual age and race groups and testing ML models that were trained on the entire dataset on these subgroups, we found that merging groups can exacerbate or reduce gaps, depending on the choice of subgroup.

For CRC, the ML models typically had higher accuracy values for the two younger age groups than the 73+ group **(Figure 2)**. This was reflected by larger percent difference values when either group’s accuracy is compared with the 73+ group’s than when the two younger groups are compared with each other **(Table 1)**. When the two higher performing groups are combined into a single subgroup, the gap in accuracy between them and the 73+ group grows for every model except MLP. However, when either group was combined with the 73+ group, the percent difference values exhibited a marked decrease, from 10.9% and 8.6% to 0.8% and 0.4% in Linear SVC **(Table 1)**.

**Table 1.**
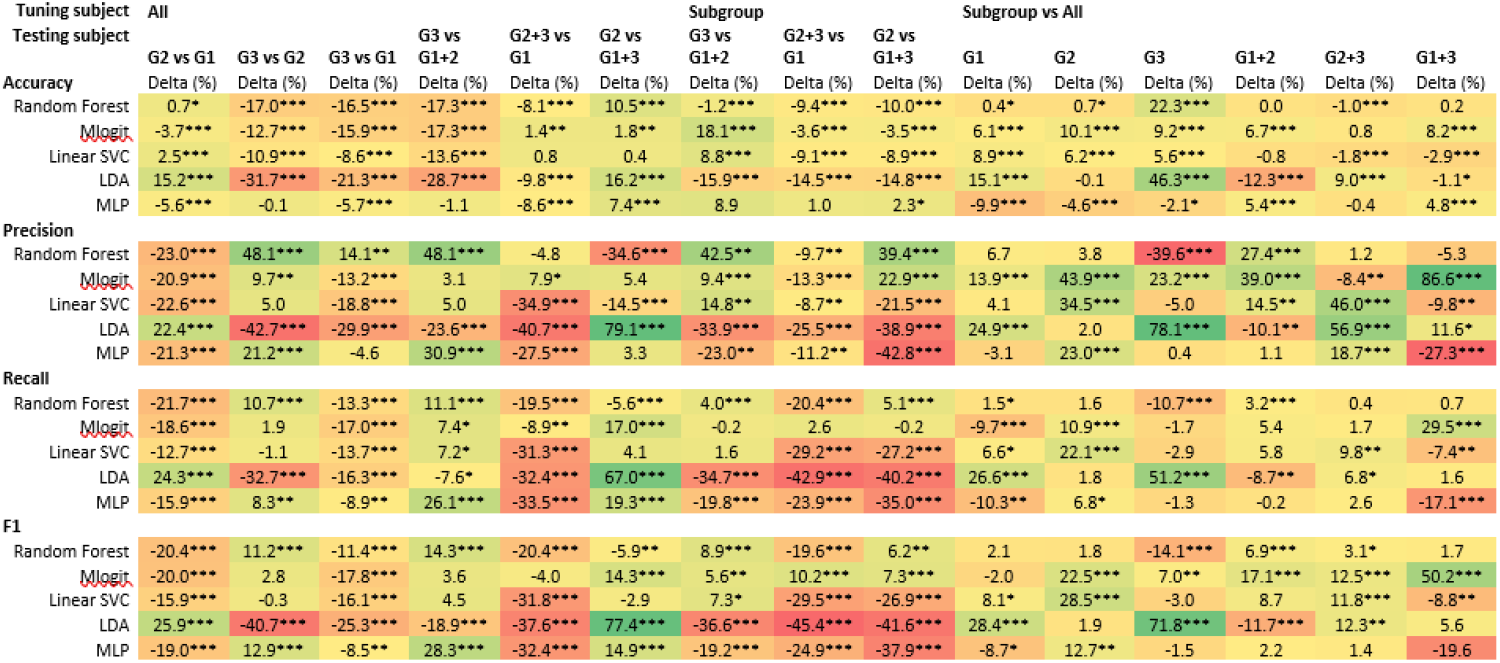
Testing whether merging CRC age groups and training the model on individual or combined groups affects the percent difference between group performance. The three groups 33-60, 61-72, and 73+ are denoted as G1, G2, and G3, respectively. The color shade indicates the difference, while the number of asterisks indicates the level of statistical differences (*, <0.05; **, <0.01; ***, <0.001).

Racial groups in CRC and Lung datasets behaved similarly. We found that we could reduce the percent difference values for performance metrics like Precision, Recall, and F1 when we combined small groups like Black patients or other minorities with a larger racial group **(Table 3 & 4)**. The improvement is most significant in MLogit, where differences in Precision as large as 73.7% and 44.2% could be reduced to 24.1% and 7.3% for CRC **(Table 3)**.

**Table 2.**
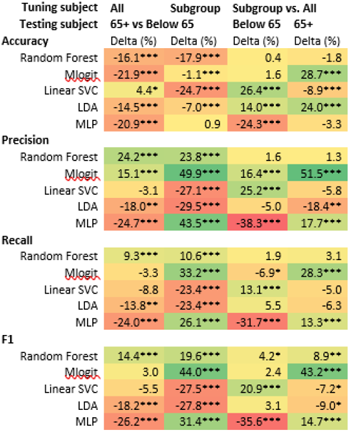
Examining the effects of ways to mitigate model bias among lung age groups. The color shade indicates the difference, while the number of asterisks indicates the level of statistical differences (*, <0.05; **, <0.01; ***, <0.001).

**Table 3.**
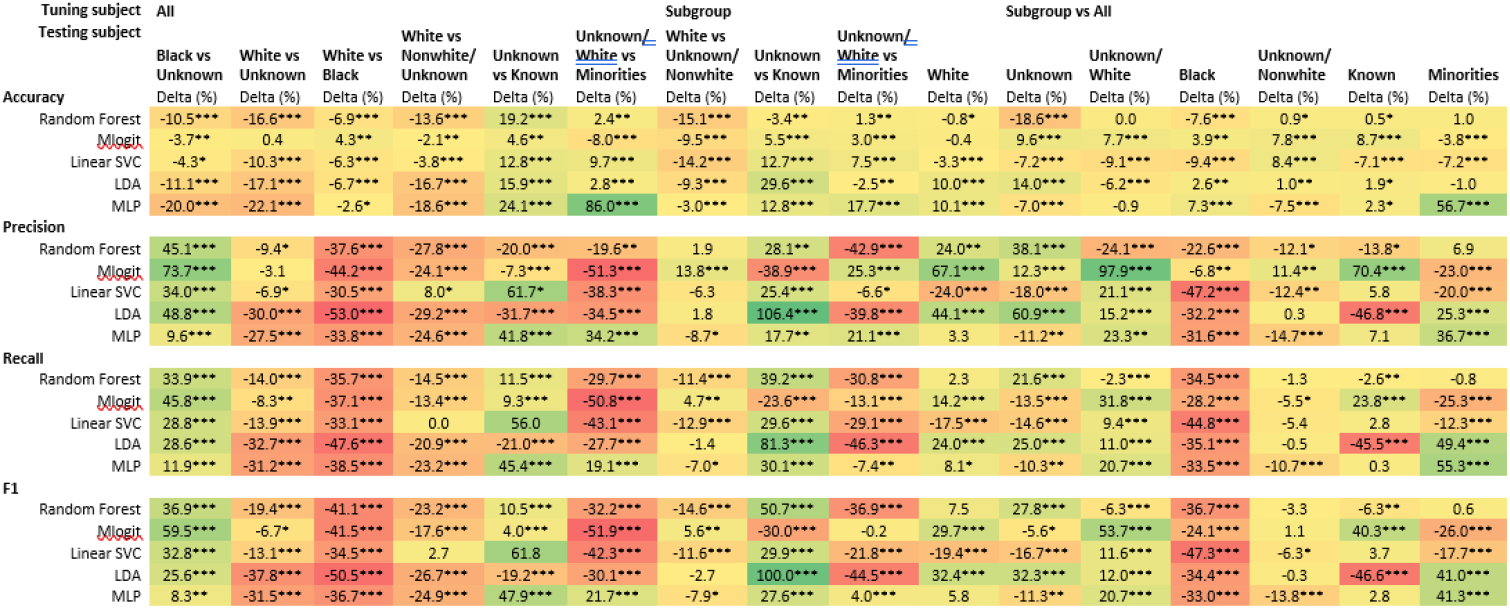
Examining the effects of ways to mitigate model bias among CRC Race groups. The color shade indicates the difference, while the number of asterisks indicates the level of statistical differences (*, <0.05; **, <0.01; ***, <0.001).

**Table 4.**
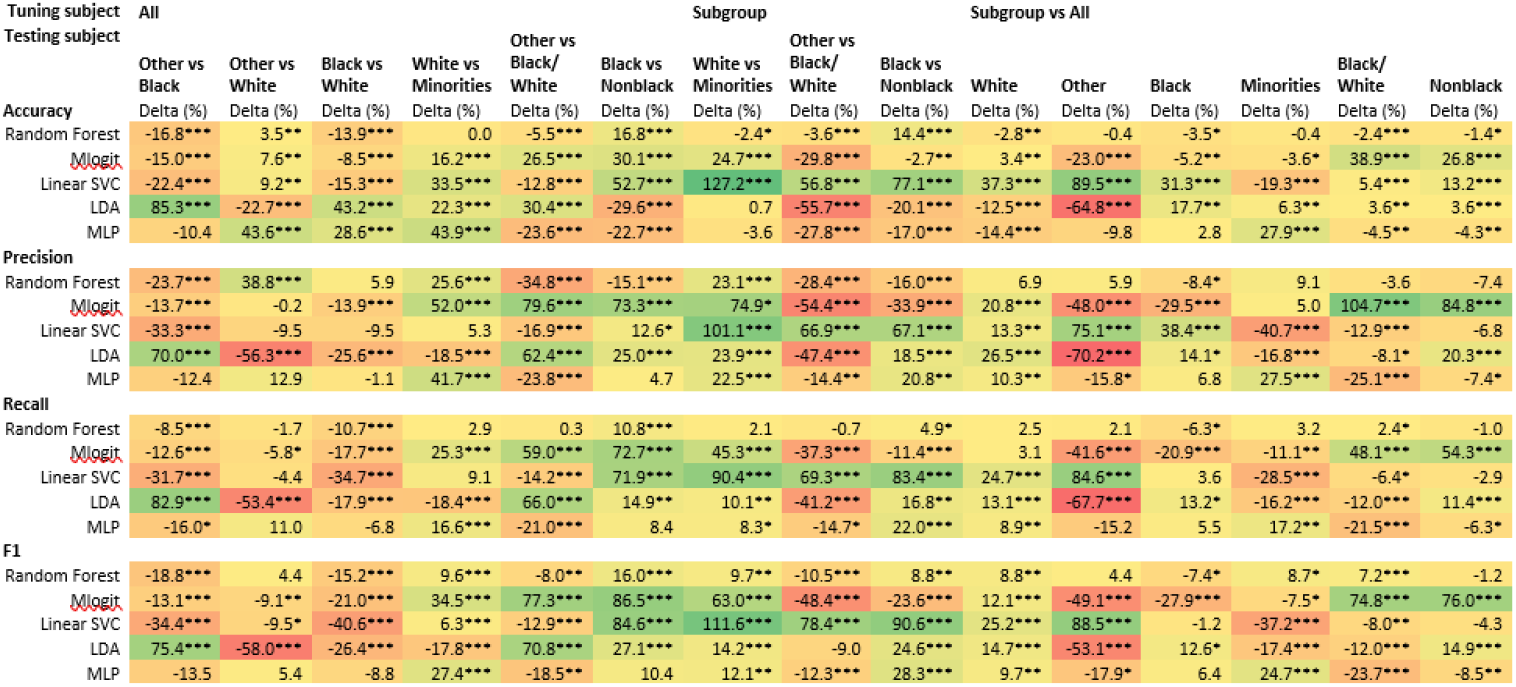
Examining the effects of ways to mitigate model bias among lung race groups. The color shade indicates the difference, while the number of asterisks indicates the level of statistical differences (*, <0.05; **, <0.01; ***, <0.001).

We also tuned the ML models on individual or combined age, sex, and race groups, before testing the model on other members of the same group to see if this could improve model performance. We found that this method seems to reduce the performance difference between the two groups for certain models without needing to penalized model performance for bias [26-28].

RF and MLP generally showed an improvement in reducing model bias when using this method. Whereas accuracy values for RF remained relatively constant for both CRC and Lung patients, Precision, Recall, and F1 typically experienced an improvement, especially among CRC Sex and Age, as well as Lung Sex **(Table 3, 5 and 6)**. Meanwhile, for MLP, this method resulted in a significant improvement among all performance metrics across many of the CRC and Lung groups, and especially for CRC and Lung Sex **(Tables 5 and 6)**.

**Table 5.**
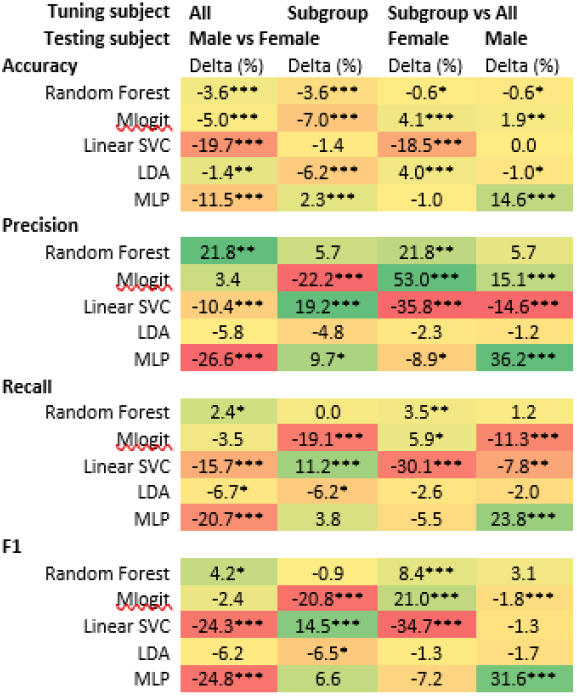
Examining the effects of ways to mitigate model bias among CRC groups based on sex. The color shade indicates the difference, while the number of asterisks indicates the level of statistical differences (*, <0.05; **, <0.01; ***, <0.001).

**Table 6.**
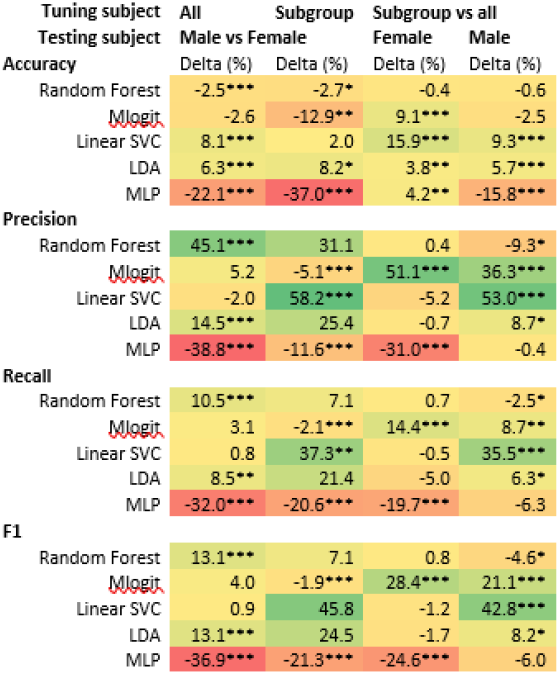
Examining the effects of ways to mitigate model bias among Lung groups based on sex. The color shade indicates the difference, while the number of asterisks indicates the level of statistical differences (*, <0.05; **, <0.01; ***, <0.001).

Linear SVC reduced the gap in accuracy values from 19.7% to 1.4% for CRC Sex and from 8.1% to 2% for Lung Sex **(Tables 5 and 6)**. However, it often increased the gap for other performance metrics, especially across the Lung sociodemographic groups (**Tables 2, 4 and 6)**.

Finally, we calculated the MSE values for each method that we used to try and reduce model bias **(Table 7)**. We found that across the same ML models and performance metrics, both merging groups and tuning and testing the model on the same group often reduced the magnitude of the MSE values. This was especially apparent in models like RF (data not shown) across both CRC and lung cancer patients when dividing them into groups based on their age, sex, and race.

**Table 7.** The mean square errors by cancer type and socioeconomical groups.

| Tuning Subject | All | All on Subgroup | Subgroup | Subgroup vs All |
| --- | --- | --- | --- | --- |
| **CRC** |  |  |  |  |
| Age | 0.0091 | 0.0110 | 0.0041 | 0.0220 |
| Gender | 0.0045 | N/A | 0.0027 | 0.0110 |
| Race | 0.0237 | 0.0216 | 0.0145 | 0.0174 |
| **Lung** |  |  |  |  |
| Age | 0.0046 | N/A | 0.0059 | 0.0071 |
| Gender | 0.0054 | N/A | 0.0086 | 0.0790 |
| Race | 0.0176 | 0.0081 | 0.0163 | 0.0246 |

## Discussion

Here we examined the impact that age, sex, and race have on the five ML models’ multimetric performance when classifying patient 4-category survival outcomes using CRC and lung adenocarcinoma data from TCGA. We confirmed that ML models exhibit significant biases and found that ML models tended to perform more poorly overall on the largest sociodemographic groups. Our data show that both merging groups (depending on the choice of subgroup) and training models on each individual or merged group hold the potential to reduce ML performance biases and thus improve AI fairness.

There are few published works evaluate ML model bias in classifying cancer outcomes using omic and clinic data, while some investigated the racial biases in ML modelling of population data[18], cancer imaging or classification[14, 20, 27, 30]. However, most, if not all, of them base evaluations of model performance solely on accuracy. This study is novel in that we compared the multimetric performance of RF, MLogit, Linear SVC, LDA, and MLP models to empirically examine model bias for different age, sex, and race groups for the first time.

Moreover, most, if not all, previous cancer studies on ML or AI fairness sort patients into binaries, especially for factors like race (comparing black and white patient groups), further marginalizing racial minorities [13, 14, 20], except one[18]. Thus, model performance and biases towards other racial groups, such as AAPI or American Natives, are poorly understood. While we were limited by the small size of these groups and were only able to examine them when merged with other groups, our findings will shed light on how model performance is impacted by algorithmic bias towards small minority groups. Furthermore, most previous ML studies on cancer survivals using transcriptomic data and socioeconomical status(s) classify patient outcomes into binary or time-based survivals [30, 31]. By analyzing four categories of outcomes, especially in the context of patient sociodemographic factors, our models can help better understand treatment outcome (e.g., a death with cancer likely linked to under-treatment) and thus inform personalized cancer treatments.

We also for the first time examined how to reduce biases toward age and gender in ML performance on classifying cancer outcomes. Several original studies focused on gender biases in ML performance[17, 23-25, 32-36], but to our knowledge few focused on cancer. Interestingly, review and guideline articles have raised the concerns on the gender biases or fairness issues[13, 37-40]. Age has been included as a covariable in several studies focused on AI/ML fairness[13, 14, 16, 18, 19, 22, 27]. However, only few focused on the fairness in age (i.e., ageism)[17]. Therefore, our works shed light on the existence and possible solutions to ML performance biases by age and may help improve ML performance in older patients who fared worse in our study.

In addition, many prior works examining AI fairness penalize the model for exhibiting bias[26-28, 41]. Despite possible overfitting, this study was able to reduce gaps in performance without doing so for RF, Linear SVC, and MLP by training models on individual or merged groups. This will allow us to reduce algorithmic bias while enabling the model to succeed when classifying patients from every group, rather than tearing down groups to equalize performance.

This study has several limitations that should be considered. We used 5-fold cross-validation and ran each model 20 times to rigorously examine the models, a method that has previously been shown to be effective [5, 9, 10]. 10-fold cross-validation may provide more samples for training, but significantly reduce the sample size in the validation/testing set. Given our focus on the minority groups, 5-fold cross-validation was chosen although 10-fold cross-validation is preferred. Moreover, the sample size of the two TCGA cohorts may be too small, especially in the context of the different sociodemographic groups. Future large-scale studies with more balanced age, sex, and race cohorts, or at least cohorts more representative of populations in the US, are warranted to better understand algorithmic bias and examine the intersectionality between different sociodemographic identities. Furthermore, we did not use the so-called fairness metrics, but the traditional group versus group comparison for robust statistical inference. The main reason is that those AI fairness metrics are mostly debatable or attributable to a single source [18, 42]. Indeed, one group appears to prefer established (balanced) accuracy as the performance metrics over fairness metrics[43]. Another group recommended using a metric best aligned with clinical purpose when assessing fairness of using ML for clinical decision support since various fairness metrics often result in conflicting results[44]. Finally, we were not able to use independent datasets that are as comprehensive as TCGA. Indeed, data sources and recruiting more patients of race-, age- and gender-minority have been recommended [45-48]. Hopefully, better and more comprehensive datasets will become available and enable us to rigorously examine our findings in an independent dataset. Additional studies are needed to validate our findings.

In summary, we here show that ML models like RF, MLogit, Linear SVC, LDA, and MLP do show bias towards different age, sex, and racial groups when classifying patients with colorectal or lung adenocarcinoma into their 4-category survival outcomes. These findings may help improve our understanding of algorithmic bias and how to include better representation of patients from all sociodemographic backgrounds. Our study also reveals several ways to design the models that hold the potential to reduce model bias.

## Supporting information

STable 1

STable 2

## Funding

This work was supported by the U.S. National Science Foundation (IIS-2128307 to LZ), the National Institute of Diabetes and Digestive and Kidney Diseases, National Institutes of Health (R01DK132885 to NG) and the National Cancer Institute, National Institutes of Health (R37CA277812 to LZ).

## Author Contributions Statement

Study conceptualization and design, ensuring the data access, accuracy and integrity (LZ), and manuscript writing (CHF and LZ). All authors, including CHF, MLD, DF, NG and LZ, contributed to the writing or revision of the review article and approved the final publication version.

## Conflicts of Interest

The authors declare no other conflict of interests.

## Data Availability Statement

The data sets used and/or analyzed of this study are available on the cBioPortal website (https://www.cbioportal.org/). The program coding is available from the corresponding authors on reasonable request.

## Compliance with ethical standards

This exempt study using publicly available de-identified data did not require an IRB review.

## Notes

### Competing Interest Statement

The authors have declared no competing interest.

