## Supplementary material for "Towards machine learning fairness in classifying multicategory causes of deaths in colorectal or lung cancer patients": STable 1

**Supplementary Table 1. Baseline characteristics of the included colorectal cancer cases by [illegible]**

| Outcome | Alive no disease<br>N (%) | Alive with disease<br>N (%) | Dead no disease<br>N (%) | Dead with disease<br>N (%) |
|---|---|---|---|---|
| **Location** | | | | |
| Colon | 293 (72) | 46 (71) | 27 (77) | 71 (84) |
| Rectal | 113 (28) | 19 (29) | 8 (23) | 14 (16) |
| **Age (yr)** | | | | |
| 33-60 | 136 (33) | 27 (42) | 1 (3) | 24 (28) |
| 61-72 | 146 (36) | 20 (31) | 11 (31) | 20 (24) |
| 73+ | 124 (31) | 18 (28) | 23 (66) | 41 (48) |
| **Sex** | | | | |
| Female | 196 (48) | 28 (43) | 22 (63) | 34 (40) |
| Male | 210 (52) | 37 (57) | 13 (37) | 51 (60) |
| **Race** | | | | |
| Unknown | 175 (43) | 17 (27) | 2 (6) | 35 (42) |
| American Native or Alaskan Indian | 1 (0) | 0 | 0 | 0 |
| Black or African American | 41 (10) | 10 (16) | 1 (3) | 12 (14) |
| White | 180 (44) | 36 (56) | 32 (91) | 35 (42) |
| Asian and Pacific Islander | 9 (2) | 1 (2) | 0 | 2 (2) |
| **Pathologic staging** | | | | |
| 1 | 90 (22) | 7 (11) | 5 (14) | 2 (2) |
| 2 | 162 (40) | 24 (37) | 12 (34) | 21 (25) |
| 3 | 112 (28) | 20 (31) | 14 (40) | 24 (28) |
| 4 | 42 (10) | 14 (22) | 4 (11) | 38 (45) |
| **Pathologic T category** | | | | |
| T1 | 18 (4) | 1 (2) | 1 (3) | 1 (1) |
| T2 | 87 (21) | 8 (12) | 6 (17) | 2 (2) |
| T3 | 269 (66) | 46 (71) | 24 (69) | 61 (72) |
| T4 | 32 (8) | 10 (15) | 4 (11) | 21 (25) |
| **Pathologic N category** | | | | |
| N0 | 262 (65) | 32 (49) | 18 (51) | 28 (33) |
| N1 | 97 (24) | 14 (22) | 7 (20) | 23 (27) |
| N2 | 46 (11) | 19 (29) | 10 (29) | 34 (40) |
| NX | 1 (0) | | | |
| **Pathologic M category** | | | | |
| M0 | 328 (82) | 45 (69) | 25 (76) | 41 (49) |
| M1 | 33 (8) | 13 (20) | 1 (3) | 36 (43) |
| MX | 41 (10) | 7 (11) | 7 (21) | 7 (8) |
| **Radiotherapy** | | | | |
| NA | 65 (16) | 2 (3) | 13 (37) | 21 (25) |
| No | 324 (80) | 54 (83) | 22 (63) | 63 (74) |

| | | | | |
|---|---|---|---|---|
| Yes | 17 (4) | 9 (14) | 0 | 1 (1) |

**outcome**
