## Supplementary material for "Towards machine learning fairness in classifying multicategory causes of deaths in colorectal or lung cancer patients": STable 2

### Supplementary Table 2. Baseline characteristics of the included lung cancer cases by outcome

| Outcome | Alive no disease, N (%) | Alive with disease, N (%) | Dead no disease, N (%) | Dead with disease, N (%) |
|---|---|---|---|---|
| **Age (yr)** | | | | |
| Below 65 | 119 (47) | 26 (34) | 30 (38) | 45 (42) |
| 65 and Above | 133 (53) | 50 (66) | 50 (63) | 62 (58) |
| **Sex** | | | | |
| Female | 140 (56) | 39 (51) | 36 (45) | 62 (58) |
| Male | 112 (44) | 37 (49) | 44 (55) | 45 (42) |
| **Race** | | | | |
| Black | 28 (11) | 7 (9) | 5 (6) | 12 (11) |
| Other | 34 (13) | 15 (20) | 17 (21) | 9 (8) |
| White | 190 (75) | 54 (71) | 58 (73) | 86 (80) |
| **Pathologic T category** | | | | |
| T1 | 107 (42) | 19 (25) | 19 (24) | 24 (22) |
| T2 | 123 (49) | 47 (62) | 44 (55) | 66 (62) |
| T3 | 15 (6) | 10 (13) | 9 (11) | 13 (12) |
| T4 | 7 (3) | 0 | 8 (10) | 4 (4) |
| **Pathologic N category** | | | | |
| N0 | 185 (73) | 57 (76) | 37 (46) | 52 (49) |
| N1 | 33 (13) | 10 (13) | 18 (23) | 35 (33) |
| N2 | 26 (10) | 7 (9) | 21 (26) | 20 (19) |
| NX | 8 (3) | 2 (3) | 4 (5) | 0 |
| **Pathologic M category** | | | | |
| M0 | 165 (63) | 48 (63) | 54 (68) | 79 (74) |
| M1 | 7 (3) | 3 (4) | 10 (13) | 5 (5) |
| MX | 80 (32) | 25 (33) | 16 (20) | 23 (21) |
| **Radiotherapy** | | | | |
| No | 79 (31) | 20 (26) | 20 (25) | 23 (21) |
| Yes | 3 (1) | 1 (1) | 2 (3) | 7 (7) |
| NA | 170 (67) | 55 (72) | 58(73) | 77 (72) |
